# Angiotensin peptides enhance SARS-CoV-2 spike protein binding to its host cell receptors

**DOI:** 10.1101/2024.12.12.628247

**Authors:** Katelin X. Oliveira, Yuichiro J. Suzuki

## Abstract

Severe Acute Respiratory Syndrome Coronavirus 2 (SARS-CoV-2), the virus that caused the Coronavirus Disease 2019 (COVID-19) pandemic, has a spike glycoprotein that is involved in recognizing and fusing to host cell receptors, such as angiotensin-converting enzyme 2 (ACE2), neuropilin-1 (NRP1), and AXL tyrosine-protein kinase. Since the major spike protein receptor is ACE2, an enzyme that regulates angiotensin II (1-8), this study tested the hypothesis that angiotensin II (1-8) influences the binding of the spike protein to its receptors. While angiotensin II (1-8) did not influence spike-ACE2 binding, we found that it significantly enhances spike-AXL binding. Our experiments showed that longer lengths of angiotensin, such as angiotensin I (1-10), did not significantly affect spike-AXL binding. In contrast, shorter lengths of angiotensin peptides, in particular, angiotensin IV (3-8), strongly increased spike-AXL binding. Angiotensin IV (3-8) also enhanced spike protein binding to ACE2 and NRP1. The discovery of the enhancing effects of angiotensin peptides on spike-host cell receptor binding may suggest that these peptides could be pharmacological targets to treat COVID-19 and post-acute sequelae of SARS-CoV-2 (PASC), which is also known as long COVID.

## Introduction

Severe Acute Respiratory Syndrome Coronavirus 2 (SARS-CoV-2) is the positive sense, single-stranded RNA virus of the genus *Betacoronavirus* that caused the Coronavirus Disease 2019 (COVID-19) pandemic [1, 2]. This virus is comprised of the following structural proteins: nucleocapsid protein, membrane protein, envelope protein, and spike protein [1]. The spike protein is the virus’s membrane fusion protein that recognizes and fuses to host cell receptors [1, 2]. The spike protein has two subunits: the S1 subunit and the S2 subunit [1, 2]. The S1 subunit contains the receptor binding domain (RBD), which directly interacts with the host cell receptor [2]. The S2 subunit anchors the spike protein to the viral membrane [1] and mediates membrane fusion [2]. After binding to the host cell receptor, the spike protein is cleaved by transmembrane protease serine 2 (TMPRSS2), activating membrane fusion and releasing the viral ribonucleoprotein complex into the host cell [2].

Angiotensin-converting enzyme (ACE) and angiotensin-converting enzyme 2 (ACE2) are key regulators of the renin-angiotensin system (RAS), which is involved in several cardiac and renal functions [3, 4]. In the classical RAS pathway, angiotensinogen, a prohormone protein, is derived in the liver and converted to angiotensin I (1-10) by renin in the kidney [3–5]. Angiotensin I (1-10) is comprised of the following ten amino acids: Asp, Arg, Val, Tyr, Ile, His, Pro, Phe, His, and Leu [4, 6]. ACE cleaves the His9 and Leu10 in angiotensin I (1-10) to produce angiotensin II (1-8) [3–6], which is comprised of Asp, Arg, Val, Tyr, Ile, His, Pro, and Phe [3, 4, 6]. Angiotensin II (1-8) binds to the type 1 angiotensin II receptor (AT_1_R), a G-protein coupled receptor (GPCR), to cause smooth muscle contraction, increase aldosterone production, antidiuretic hormone (ADH) production, sympathetic nervous system tone, blood pressure, cardiac hypertrophy, and fibrosis, and decrease nitric oxide (NO) production, parasympathetic nervous system tone, baroreflex sensitivity, and natriuresis [4, 6, 7]. Angiotensin II (1-8) also binds to another GPCR called the type 2 angiotensin II receptor (AT_2_R), which mediates opposing effects of angiotensin II (1-8) via the AT_1_R [3, 5, 7]. The actions of AT_2_R have vasodilatory, anti-fibrotic, anti-inflammatory, and anti-proliferative properties [3–5, 7]. Additionally, angiotensin II (1-8) stimulates reactive oxygen species production, cell growth, cell differentiation, apoptosis, and inflammatory responses [5, 7]. Aminopeptidase A (APA) then cleaves the Asp1 on angiotensin II (1-8), producing angiotensin III (2-8) [3–5, 7]. Angiotensin III (2-8) also functions through AT_1_R and AT_2_R [4, 5], increasing blood pressure and ADH release, but with a lower potency [4, 5, 7]. Following, aminopeptidase N (APN) converts angiotensin III (2-8) into angiotensin IV (3-8) by removing Arg2 [3–5]. Angiotensin IV (3-8) binds to the type 4 angiotensin II receptor (AT_4_R) [3, 4, 5], which is expressed in the brain, kidney, heart, and vessels [5], to increase natriuresis and NO production and stimulate cardioprotective effects, such as reduced vasoconstriction [4]. In addition to contributing to blood flow regulation [3, 5], angiotensin IV (3-8) is also involved in learning [3, 8], memory [3, 4, 8], and neuronal development [3, 8]. In the counter-regulatory RAS pathway, ACE2 can generate angiotensin (1-9) from angiotensin I (1-10) by removing Leu10 [4, 6]. Angiotensin (1-9) exerts its effects through AT_2_R to activate natriuresis and NO production [4]. Additionally, angiotensin I (1-10) can be cleaved by neprilysin (NEP) to produce angiotensin (1-7) [4, 6]. Furthermore, ACE2 can cleave off the Phe8 in angiotensin II (1-8), a potent vasoconstrictor, to produce angiotensin (1-7), a vasodilator [3, 4]. Angiotensin (1-7) binds to Mas, a GPCR [3, 6]. See Figure 1 for schematics of angiotensin metabolism.

**Figure 1:**
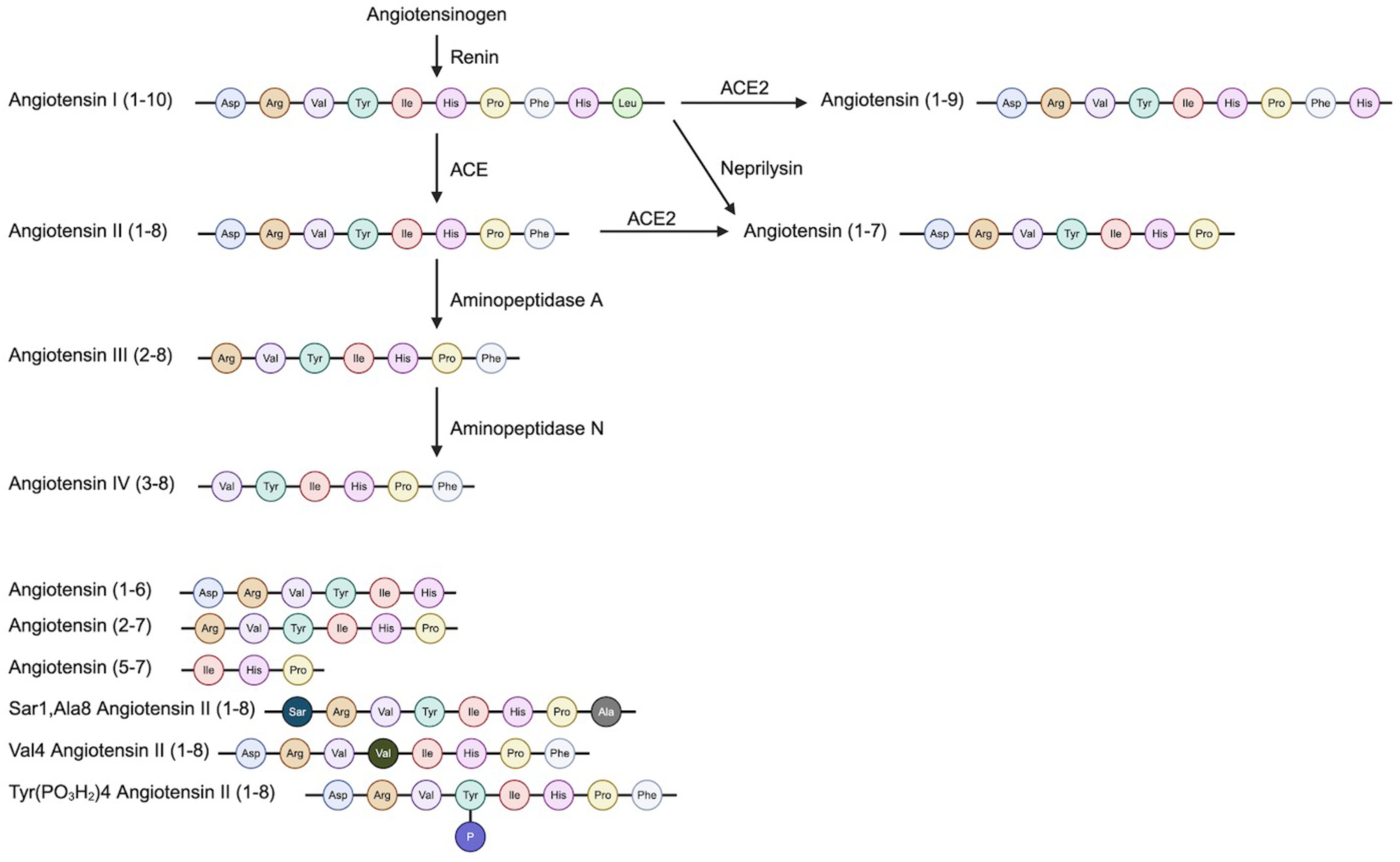
Structures, Synthesis, and Metabolism of the Angiotensin peptides. This figure illustrates the structures of the angiotensin peptides relevant to this series of experiments. It also shows how angiotensin I (1-10), angiotensin (1-9), angiotensin II (1-8), angiotensin (1-7), angiotensin III (2-8), and angiotensin IV (3-8) are physiologically synthesized and metabolized in the human body.

In addition to playing an essential role in hemodynamics and regulating angiotensin II (1-8) levels, it has been well established that ACE2 acts as a receptor for SARS-CoV-2 [1, 2, 9, 10]. The SARS-CoV-2 infection of human airway epithelia correlates with well-differentiated cells that highly express ACE2 [11]. Although ACE2 is the major receptor for SARS-CoV-2 entry, ACE2 expression is low in the respiratory system [12–14], particularly in pulmonary and bronchial tissue, suggesting that other receptors are needed to facilitate SARS-CoV-2 entry [14]. ACE2 expression was mainly found in enterocytes, gallbladder, male reproductive cells, placental trophoblasts, cardiomyocytes, vasculature, eye, and kidneys [12, 14].

A candidate receptor for SARS-CoV-2 is AXL [14]. AXL is a receptor tyrosine kinase [14–18] and a member of the TAM family, which includes TYRO_3_, AXL, and MER [14–16]. This receptor tyrosine kinase mediates signaling pathways and is associated with various biological processes, including proliferation, migration, differentiation, and apoptosis [15, 16]. Mainly involved in cancer research, AXL was discovered in patients with chronic myeloid leukemia [15–19]. AXL expression is observed in vascular cells, immune cells [15], embryonic cells [16], tumor cells, bone marrow stoma [17], fibroblasts, myeloid progenitor cells, neural tissue, cardiac muscle, and skeletal muscle [18]. Notably, a higher percentage of lung and tracheal cells express AXL [14]. Additionally, previous studies suggest that AXL expression may be upregulated by G protein-coupled receptor agonists, such as angiotensin II (1-8) and thrombin [19]. In addition to being a receptor for SARS-CoV-2, AXL is a potential molecular marker for COVID-19 progression [20].

In the present study, we aimed to examine the effects of angiotensin II (1-8) on spike protein binding to host cell receptors. We found that although spike-ACE2 binding was not modified by angiotensin II (1-8), angiotensin II (1-8) enhanced spike-AXL binding. We further investigated the effects of other angiotensins on spike-AXL binding.

## Materials and Methods

### Chemicals

Angiotensin I (1-10), angiotensin (1-9), angiotensin II (1-8), angiotensin (1-7), angiotensin (1-6), angiotensin III (2-8), angiotensin (2-7), and angiotensin (5-7) were purchased from APExBIO Technology LLC (Houston, TX, USA). Angiotensin IV (3-8) was purchased from Bachem Americas, Inc. (Torrance, CA, USA). Sar1, Ala8 angiotensin II (1-8), Val4 angiotensin II (1-8), and Tyr(PO_3_H_2_)4 angiotensin II (1-8) were purchased from MilliporeSigma (Burlington, MA, USA). Recombinant human ACE2 was purchased from Sino Biological, Inc. (Wayne, PA, USA). See Figure 1 for amino acid sequences of angiotensin peptides used in this study.

### Spike protein binding assays

RayBio COVID-19 Spike-AXL Binding Assay Kits (Catalog #: CoV-AXLS1-1) and RayBio COVID-19 Spike-NRP1 binding Assay Kits (Catalog #: CoV-NRP1S1-1) were purchased from RayBiotech Life, Inc. (Peachtree Corners, GA, USA). CoviDrop SARS-CoV-2 Spike-ACE2 Binding Inhibitor Screening Fast Kits (Catalog #: D-1004-96) were purchased from Epigentek Group Inc (Farmingdale, NY, USA). Assays were conducted according to the manufacturers’ instructions and the horseradish peroxidase (HRP) activity was measured at the absorbance of 450 nm using an EMax Plus Microplate Reader (Molecular Devices, LLC, San Jose, CA, USA).

### Statistical Analysis

The mean absorbance values and standard errors of the mean (SEM) were calculated. The control groups without angiotensin and experimental groups with angiotensin were compared using an unpaired t-test. Statistical significance was determined by a p-value of less than 0.05. To compare across different lengths of angiotensin and different spike protein receptors, the mean absorbance values were normalized to percent absorbances.

## Results

### The effects of angiotensin II (1-8) on spike protein binding

The RayBio COVID-19 Spike-AXL Binding Assay Kits were used to explore spike-AXL binding. These kits had AXL protein coated at the bottom of the well plate. Spike protein was then added to the wells and adhered to the AXL protein. After washing unbound components, the anti-spike protein antibody was applied followed by an HRP-conjugated secondary antibody to detect spike-AXL binding. To test the effect of angiotensin II (1-8) on spike-AXL binding, 40 μM of angiotensin II (1-8) were applied to the assay kits when the spike protein was added. To account for the differences in the untreated, spike-protein-only control groups over separate experiments, percent absorbance was determined. The control group’s mean percent absorbance was 100% and the angiotensin II (1-8) group’s mean percent absorbance was 200.9%, suggesting that the addition of the angiotensin II (1-8) increased spike-AXL binding. The calculated p-value was less than 0.0001, indicating that the difference between the control group and the angiotensin II (1-8) group was statistically significant (Fig. 2A).

**Figure 2:**
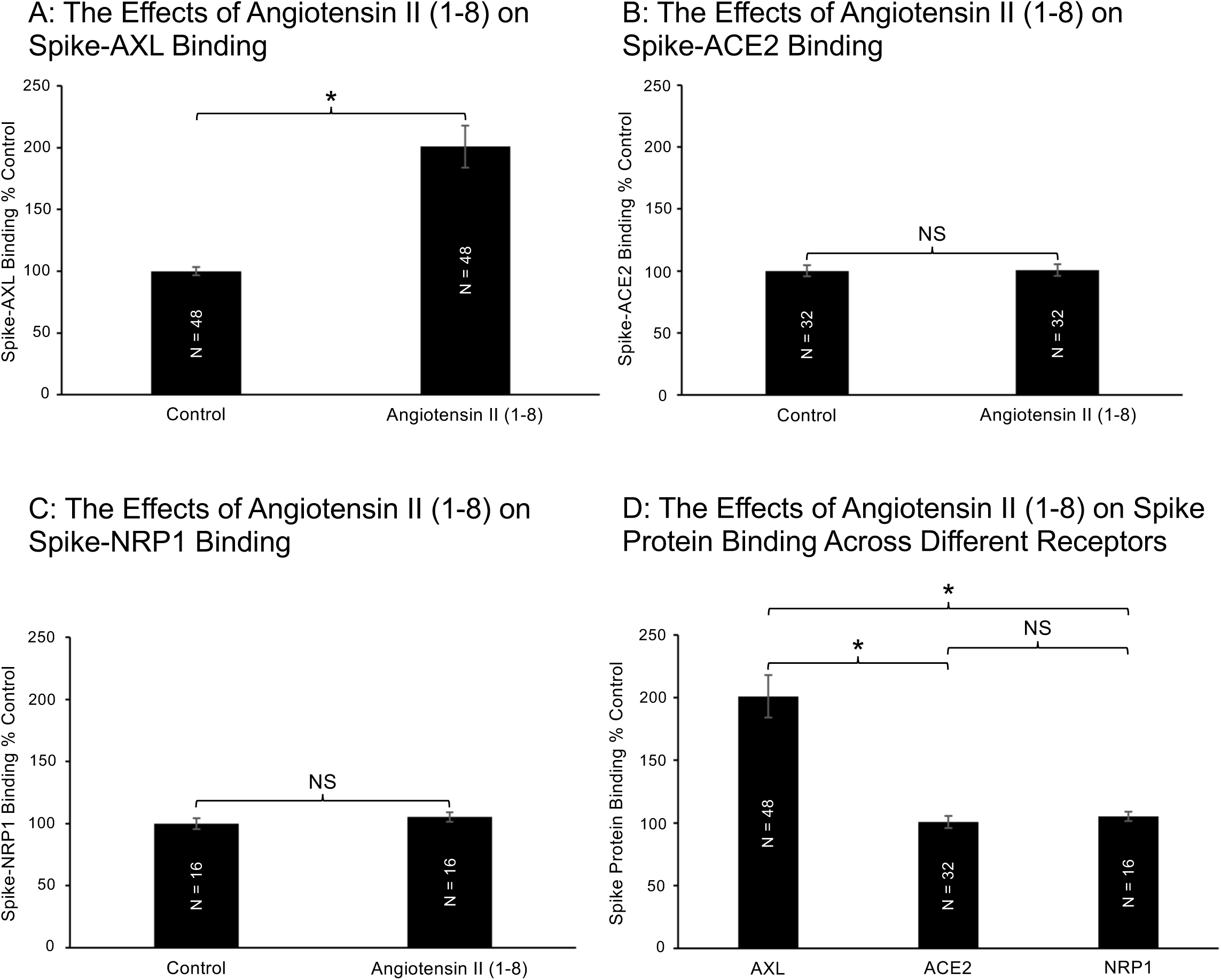
The Effects of Angiotensin II (1-8) on Spike Protein Binding. This figure depicts the effect of angiotensin II (1-8) on spike protein binding. In Figure 2A, the addition of angiotensin II (1-8) significantly enhanced spike-AXL binding. In contrast, Figures 2B and 2C show that angiotensin II (1-8) did not significantly alter spike-ACE2 binding and spike-NRP1 binding, respectively. Figure 2D combines the data in Figures 2A, 2B, and 2C to show the effect of angiotensin II (1-8) on spike protein binding across different receptors. Angiotensin II (1-8) selectively enhanced spike-AXL binding and did not influence spike-ACE2 binding and spike-NRP1 binding. One asterisk (*) denotes a p-value equal to or less than 0.05 and “NS” represents “not significant”.

CoviDrop SARS-CoV-2 Spike-ACE2 Binding Inhibitor Screening Fast Kits were used to understand the binding interactions between the spike protein and ACE2. Spike protein was coated on the bottom of the well plate and His-ACE2 was added to interact with the spike protein. After washing, anti-His antibody-HRP and binding detection solution were applied to detect spike-ACE2 binding. To assess the effect of angiotensin II (1-8) on spike-ACE2 binding, 40 μM of angiotensin II (1-8) were applied to the assay kits when His-ACE2 was added. The angiotensin II (1-8) group’s mean percent absorbance was 100.7%. The difference between the control group’s mean percent absorbance of 100% and the angiotensin II (1-8) group was not statistically significant with a p-value of 0.925, suggesting that the addition of angiotensin II (1-8) did not significantly modify spike-ACE2 binding (Fig. 2B).

To further ensure that the enhancing effect of angiotensin II (1-8) is specific to spike-AXL binding, we examined the effect of angiotensin II (1-8) on spike-NRP1 binding using RayBio COVID-19 Spike-NRP1 binding Assay Kits. Like the spike-AXL assay kits, the spike-NRP1 kits have NRP1 receptors coated on the well plate, and spike protein is added so that it attaches to the NRP1 receptors. Following the washing of unbound components, the anti-spike protein antibody and HRP-conjugated secondary antibody were added. Angiotensin II (1-8) was added to the kits as the spike protein was added to investigate how angiotensin II (1-8) modifies spike-NRP1 binding. The control group’s mean percent absorbance was 100%. The mean percent absorbance of the angiotensin II (1-8) group was 105.2%. The calculated p-value was 0.3844, suggesting that the addition of angiotensin II (1-8) did not significantly influence spike-NRP1 binding (Fig. 2C).

The use of the three binding assay kits demonstrated that the enhancing effect of angiotensin II (1-8) is specific to spike-AXL binding since angiotensin II (1-8) did not modify spike-ACE2 and spike-NRP1 binding. The difference between the effect of angiotensin II (1-8) on spike-AXL binding and the effect of angiotensin II (1-8) on spike-ACE2 binding was determined to be statistically significant due to an unpaired t-test calculating a p-value of less than 0.0001. Similarly, an unpaired t-test found that the difference between the effect of angiotensin II (1-8) on spike-AXL binding and the effect of angiotensin II (1-8) on spike-NRP1 binding was statistically significant with a p-value of 0.0021. In contrast, the difference between the effect of angiotensin II (1-8) on spike-ACE2 binding and the effect of angiotensin II (1-8) on spike-NRP1 binding was insignificant with a p-value of 0.5392. This further illustrated that the enhancing effect of angiotensin II (1-8) was specific to spike-AXL binding (Fig. 2D).

### The effects of angiotensin (1-9) and angiotensin I (1-10) on spike protein binding

Wanting to explore whether longer angiotensins affect spike-AXL binding, angiotensin (1-9) and angiotensin I (1-10) were applied to the spike-AXL assay kits. The control group’s mean percent absorbance during this experiment was 100%. The addition of angiotensin II (1-8) increased the mean percent absorbance of the angiotensin II (1-8) group to 149.9%. The increase due to the addition of angiotensin II (1-8) was found to be statistically significant after an unpaired t-test calculated a p-value of 0.0005. When 40 μM of angiotensin (1-9) were added to the kit, the mean percent absorbance of the experimental group was 113.8%. An unpaired t-test determined that the difference between the angiotensin (1-9) and the control groups was 0.3519, indicating that the addition of angiotensin (1-9) did not significantly modify spike-AXL binding. When comparing the angiotensin II (1-8) group and the angiotensin (1-9) group, the unpaired t-test determined the difference between the groups to be statistically significant with a p-value of 0.0489. Therefore, the addition of angiotensin II (1-8) significantly enhanced spike-AXL binding when compared to the control group and the angiotensin (1-9) group (Fig. 3A).

**Figure 3:**
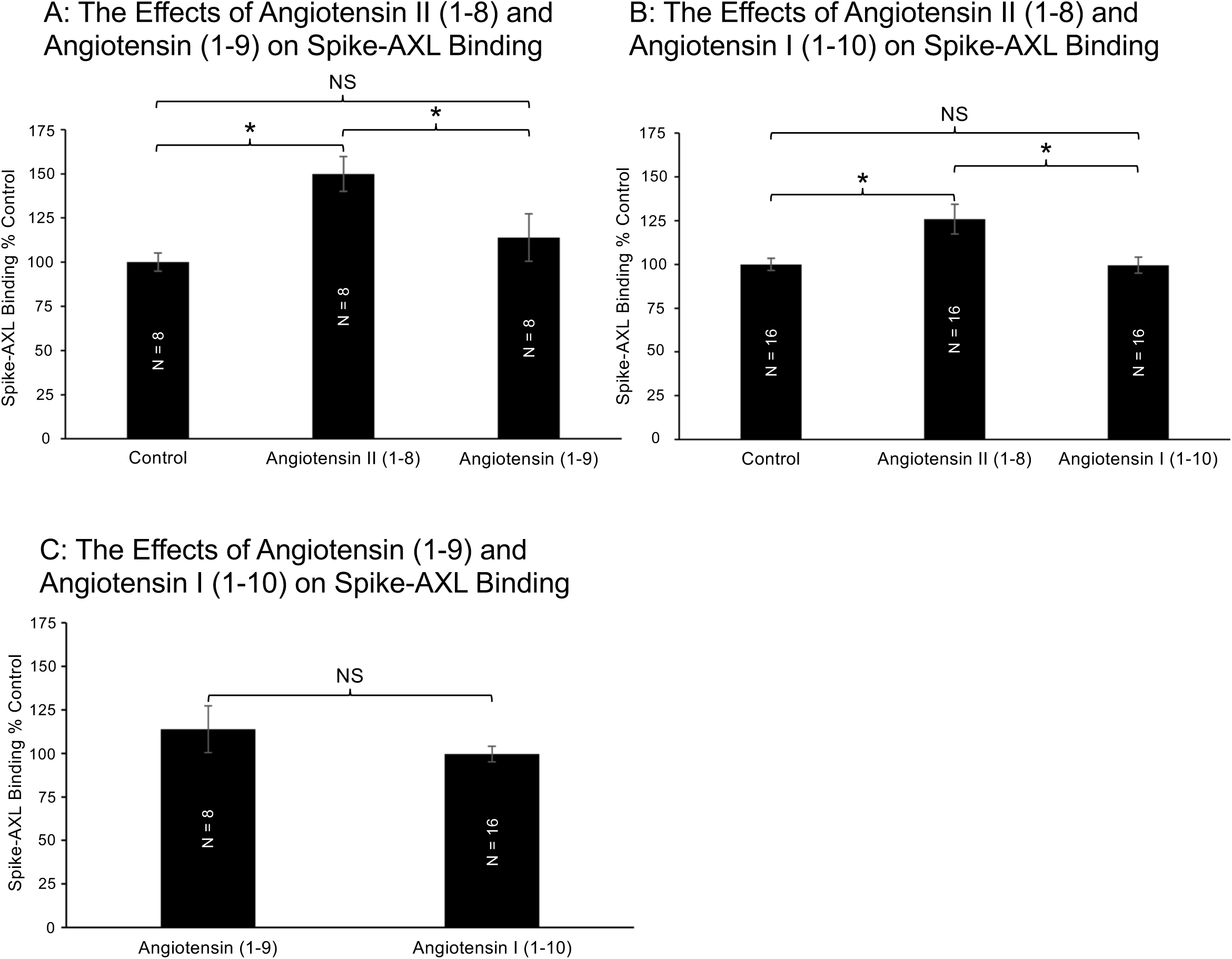
The Effects of Angiotensin II (1-8), Angiotensin (1-9), and Angiotensin I (1-10) on Spike-AXL Binding. This figure showcases the effect of angiotensin II (1-8), angiotensin (1-9), and angiotensin I (1-10) on spike-AXL binding. In Figure 3A, only angiotensin II (1-8) enhances binding, while angiotensin (1-9) does not significantly increase binding. Additionally, the difference between the effect of angiotensin II (1-8) and the effect of angiotensin (1-9) is significant, further suggesting that angiotensin II (1-8) has an enhancing effect on spike-AXL. Similarly, in Figure 3B, only angiotensin II (1-8) increases spike-AXL binding, while angiotensin I (1-10) does not. Figure 3C combines the data shown in Figures 3A and 3B to demonstrate that the difference between the effect of angiotensin (1-9) and the effect of angiotensin I (1-10) on spike-AXL binding is not significant. One asterisk (*) denotes a p-value equal to or less than 0.05 and “NS” represents “not significant”.

During this experiment, the control group’s mean percent absorbance was 100%. The addition of angiotensin II (1-8) increased the mean percent absorbance to 125.9%. The increase of the mean percent absorbance due to the addition of angiotensin II (1-8) was determined to be statistically significant by an unpair t-test producing a p-value of 0.0086. Similar to angiotensin (1-9), when 40 μM of angiotensin I (1-10) were applied to the spike-AXL assay kit, the mean percent absorbance of the angiotensin I (1-10) group was 99.6%. An unpaired t-test determined that the difference between the angiotensin I (1-10) and the control groups was not statistically significant with p-values of 0.9479. This finding suggests that the addition of angiotensin I (1-10) did not significantly modify spike-AXL binding. The difference between the angiotensin II (1-8) group and the angiotensin I (1-10) was statistically significant with a p-value of 0.0107. When compared to the control group and the angiotensin I (1-10), the addition of angiotensin II (1-8) significantly increased spike-AXL binding (Fig. 3B).

The difference between the effect of angiotensin (1-9) on spike-AXL binding and the effect of angiotensin I (1-10) on spike-AXL binding was determined to not be statistically significant due to an unpaired t-test calculating a p-value of less than 0.2208. Therefore, these experiments illustrated that angiotensin II (1-8) has an enhancing effect on spike-AXL binding, while longer angiotensins, such as angiotensin (1-9) and angiotensin I (1-10), do not influence spike-AXL binding. Angiotensin I (1-10), a precursor to angiotensin II (1-8), did not modify spike-AXL binding, suggesting that the loss of His9 and Leu10 altered the amino acid’s effect, which resulted in angiotensin II (1-8)’s enhancing effect on spike-AXL binding (Fig. 3C).

### The effects of angiotensin (1-7) and angiotensin (1-6) on spike-AXL binding

To better understand the effects of carboxyl-terminal (C-terminal) deletion from angiotensin II (1-8) on spike-AXL binding, 40 μM of angiotensin (1-7) and angiotensin (1-6) were applied to the spike-AXL assay kits. The control group’s mean percent absorbance was 100%. Adding angiotensin II (1-8) increased the mean percent absorbance to 290.3%. An unpaired t-test found that the addition of angiotensin II (1-8) significantly enhanced spike-AXL binding with a p-value of less than 0.0001. The mean percent absorbances of the angiotensin (1-7) and angiotensin (1-6) groups increased to 277.5% and 341.8%, respectively. Similar to adding angiotensin II (1-8), the enhancing effects of angiotensin (1-7) and angiotensin (1-6) were determined to be significant with p-values of less than 0.0001. To compare the mean percent absorbances between the angiotensin II (1-8), angiotensin (1-7), and angiotensin (1-6) groups, unpaired t-tests were performed. The differences between the angiotensin II (1-8) and angiotensin (1-7) groups, angiotensin II (1-8) and angiotensin (1-6) groups, and the angiotensin (1-7) and angiotensin (1-6) groups were not statistically significant with p-values of 0.8129, 0.4205, and 0.2985, respectively. The results of this experiment suggested that the C-terminal deletion of Phe8 and Pro7 from angiotensin II (1-8) did not modify the enhancing effect on spike-AXL binding (Fig. 4).

**Figure 4:**
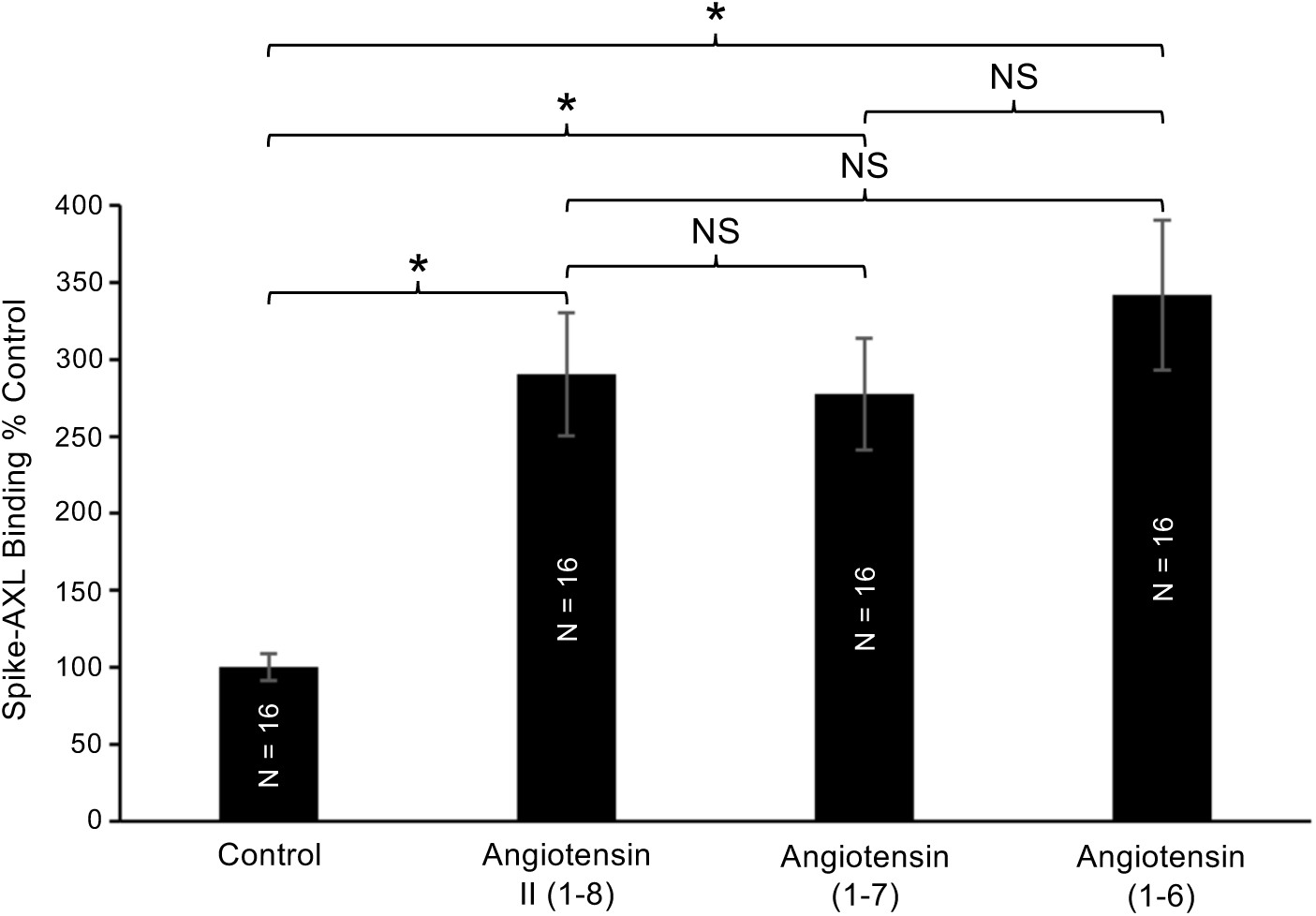
The Effects of Angiotensin II (1-8), Angiotensin (1-7), and Angiotensin (1-6) on Spike-AXL Binding. This figure illustrates how angiotensin II (1-8), angiotensin (1-7), and angiotensin (1-6) significantly enhance spike-AXL binding. However, when compared to each other, the enhancing effects of angiotensin II (1-8), angiotensin (1-7), and angiotensin (1-6) are not significantly different from one another. One asterisk (*) denotes a p-value equal to or less than 0.05 and “NS” represents “not significant”.

### The effects of angiotensin III (2-8) and angiotensin IV (3-8) on spike-AXL binding

To study the effects of amino-terminal (N-terminal) deletion from angiotensin II (1-8) on spike-AXL binding, 40 μM of angiotensin III (2-8) and angiotensin IV (3-8) were applied to the spike-AXL assay kits. The control group’s mean percent absorbance was 100%. When angiotensin II (1-8) was applied to the spike-AXL kit, the mean percent absorbance was 125.9%. The unpaired t-test that compared the control and angiotensin II (1-8) groups yielded a p-value of 0.0086, demonstrating statistical significance. Adding angiotensin III (2-8) enhanced the mean percent absorbance to 173.7%. When comparing the angiotensin III (2-8) and control groups, the difference was determined to be statistically significant with a p-value of 0.0002. Similarly, the unpaired t-test that compared the angiotensin II (1-8) and angiotensin III (2-8) groups calculated a p-value of 0.0162, indicating significance. This enhancing effect of angiotensin III (2-8) on spike-AXL binding suggests that the loss of Asp1 not only had no effect on angiotensin II (1-8)’s enhancing effect on spike-AXL binding but also increased the enhancing effect (Fig. 5A).

**Figure 5:**
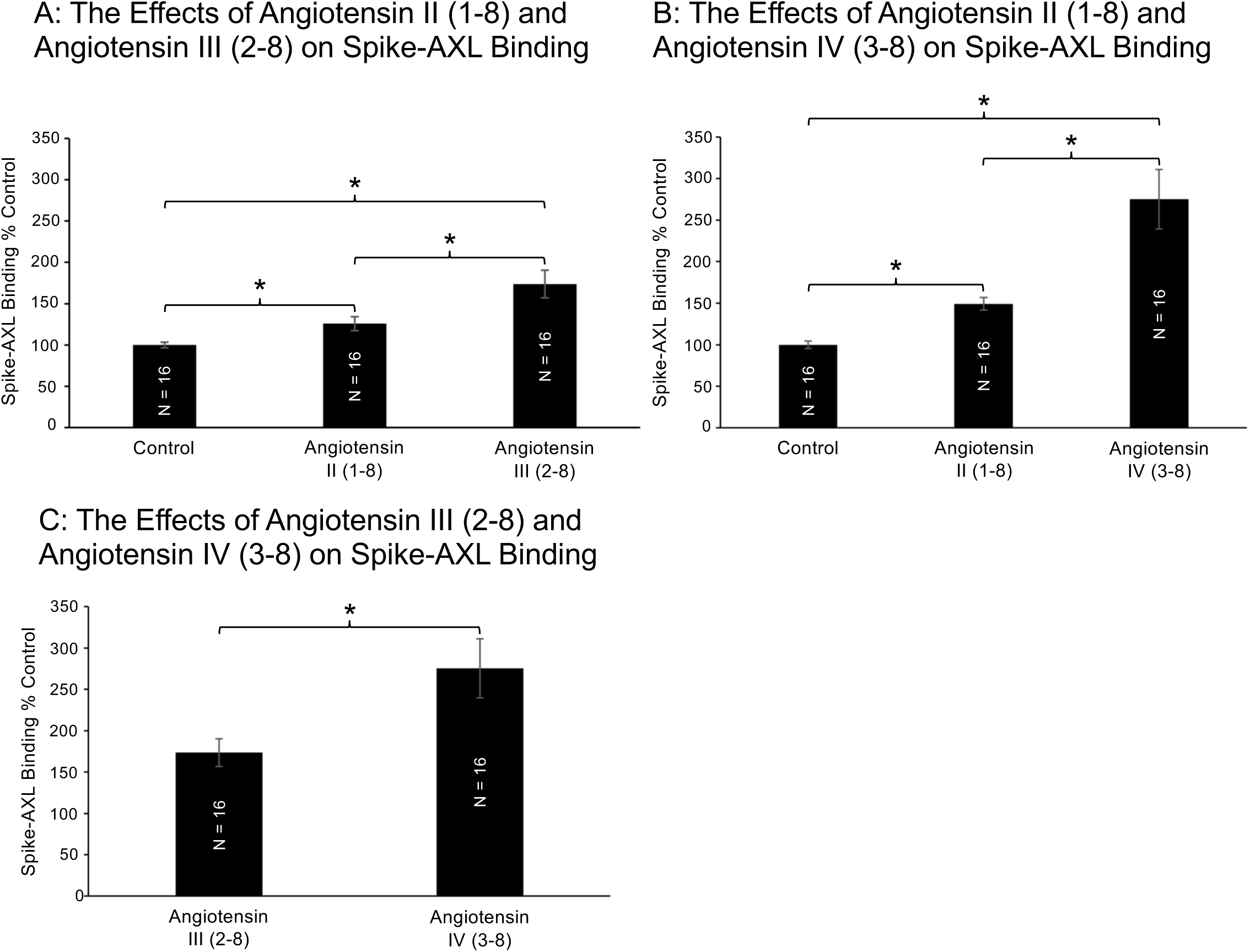
The Effects of Angiotensin II (1-8), Angiotensin III (2-8), and Angiotensin IV (3-8) on Spike-AXL Binding. This figure illustrates the effects of angiotensin II (1-8), angiotensin III (2-8), and angiotensin IV (3-8) on spike-AXL binding. In Figure 5A, both angiotensin II (1-8) and angiotensin III (2-8) enhance spike-AXL binding. Additionally, the difference between the effect of angiotensin II (1-8) and the effect of angiotensin III (2-8) is significant, suggesting that angiotensin III (2-8) enhances spike-AXL binding significantly more than angiotensin II (1-8). Similarly, Figure 5B shows that both angiotensin II (1-8) and angiotensin IV (3-8) significantly increase spike-AXL binding. There is a significant difference between the effect of angiotensin II (1-8) and the effect of angiotensin IV (3-8), indicating that angiotensin IV (3-8)’s enhancing effect on spike-AXL binding is greater than angiotensin II (1-8)’s enhancing effect. Figure 5C combines the data shown in Figures 5A and 5B to demonstrate that angiotensin IV (3-8)’s enhancement of spike-AXL binding is significantly greater than angiotensin III (2-8)’s enhancement of spike-AXL binding. One asterisk (*) denotes a p-value equal to or less than 0.05 and “NS” represents “not significant”.

To assess the importance of removing Asp1 and Arg2 in regards to N-terminal deletion from angiotensin II (1-8) on spike-AXL binding, 40 μM of angiotensin IV (3-8) were applied to the spike-AXL assay kits. The mean percent absorbance of the control group was 100%. The mean percent absorbance of the group that contained angiotensin II (1-8) was 149.2%. An unpaired t-test found that the difference between the control and angiotensin II (1-8) groups was statistically significant with a p-value of 0.0086. The addition of angiotensin IV (3-8) increased the mean percent absorbance to 275.3%. The enhancement of spike-AXL binding by angiotensin IV (3-8) was found to be statistically significant when compared to the control group with a p-value of less than 0.0001. When comparing the angiotensin II (1-8) and angiotensin IV (3-8) groups, an unpaired t-test found that the difference was statistically significant with a p-value of 0.0017, illustrating that the removal of Asp1 and Arg2 from angiotensin II (1-8) to generate angiotensin IV (3-8) greatly enhanced spike-AXL binding (Fig. 5B).

The difference between the effect of angiotensin III (2-8) on spike-AXL binding and the effect of angiotensin IV (3-8) on spike-AXL binding was determined to be statistically significant with an unpaired t-test calculating a p-value of 0.0151. This comparison suggests that the removal of Arg2 from angiotensin III (2-8) to create angiotensin IV (3-8) is essential to the further enhancement of spike-AXL binding. Therefore, these series of experiments demonstrated that the addition of angiotensin II (1-8) enhanced spike-AXL binding in comparison to the control. Additionally, the removal of Asp1 from angiotensin II (1-8) to produce angiotensin III (2-8) further strengthened spike-AXL binding when compared to the control and angiotensin II (1-8) groups. Lastly, the deletion of Arg2 from angiotensin III (2-8) to generate angiotensin IV (3-8) greatly enhanced spike-AXL binding in comparison to the control, angiotensin II (1-8), and angiotensin III (2-8) groups (Fig. 5C).

### The effects of angiotensin IV (3-8) on spike protein binding

As shown in Figure 5B, adding angiotensin IV (3-8) to the spike-AXL assay kits drastically increased the mean percent absorbance to 275.3%. When compared to the control group’s mean percent absorbance of 100%, angiotensin IV (3-8)’s enhancing effect was determined to be statistically significant with a p-value of less than 0.0001 (Fig. 6A).

**Figure 6:**
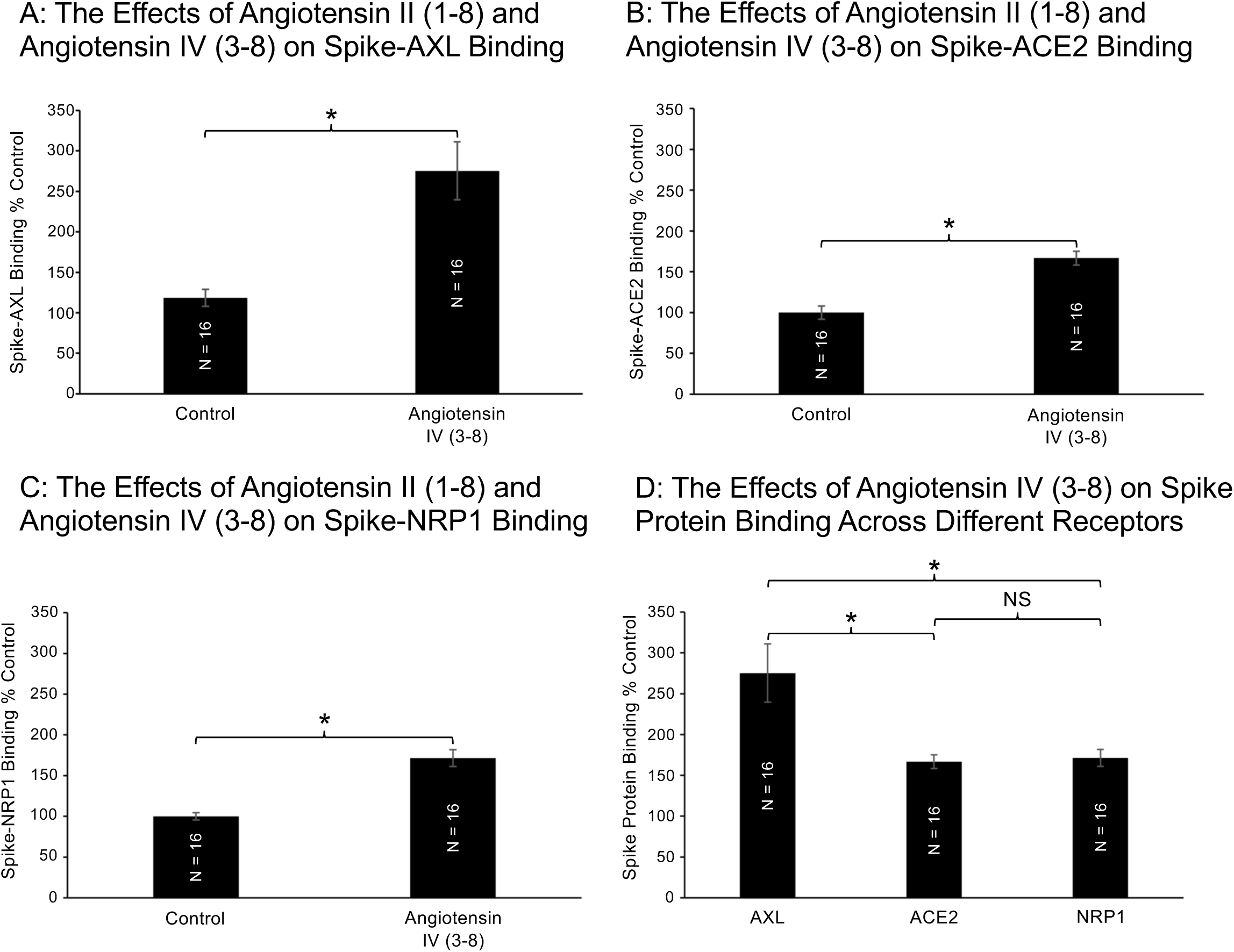
The Effects of Angiotensin IV (3-8) on Spike Protein Binding. This figure presents the effect of angiotensin IV (3-8) on spike protein binding. Figures 6A, 6B, and 6C all show that adding angiotensin IV (3-8) significantly enhanced spike-AXL binding, spike-ACE2 binding, and spike-NRP1 binding, respectively. Figure 6D combines the data in Figures 6A, 6B, and 6C to illustrate the effect of angiotensin IV (3-8) on spike protein binding across different receptors. Although angiotensin IV (3-8) enhanced spike-AXL binding, spike-ACE2 binding, and spike-NRP1 binding, the difference angiotensin IV (3-8)’s enhancement of spike-AXL binding and angiotensin IV (3-8)’s enhancement of spike-ACE2 binding and the difference angiotensin IV (3-8)’s enhancement of spike-AXL binding and angiotensin IV (3-8)’s enhancement of spike-NRP1 binding are significant, implying that angiotensin IV (3-8) enhanced spike-AXL binding more than it increased spike-ACE2 binding and spike-NRP1 binding. One asterisk (*) denotes a p-value equal to or less than 0.05 and “NS” represents “not significant”.

After discovering angiotensin IV (3-8)’s enhancing effect on spike-AXL binding, angiotensin IV (3-8)’s effect on spike-ACE2 binding and spike-NRP1 binding was studied. To assess the effect of angiotensin IV (3-8) on spike-ACE2 binding, 40 μM of angiotensin IV (3-8) were applied to the spike-ACE2 assay kits. Applying angiotensin IV (3-8) increased the mean percent absorbance to 166.7%. The difference between the control group’s mean percent absorbance of 100% and the angiotensin IV (3-8) group was statistically significant with a p-value of less than 0.0001, suggesting that angiotensin IV (1-8) had an enhancing effect on spike-ACE2 binding (Fig. 6B).

Similarly, 40 μM of angiotensin IV (3-8) were applied to the spike-NRP1 assay kits. The control group’s mean percent absorbance was 100%. The mean percent absorbance of the angiotensin IV (3-8) group was 171.4%. In comparison to the control group, the increase of the mean percent absorbance due to the addition of angiotensin IV (3-8) was determined to be significant with a p-value of less than 0.0001 (Fig. 6C).

Like spike-AXL binding, angiotensin IV (3-8) has enhancing effects on spike-ACE2 binding and spike-NRP1 binding. The difference between the effect of angiotensin IV (3-8) on spike-AXL binding and the effect of angiotensin IV (3-8) on spike-ACE2 binding was found to be statistically significant with a p-value of 0.0060. Similarly, the difference between the effect of angiotensin IV (3-8) on spike-AXL binding and the effect of angiotensin IV (3-8) on spike-NRP1 binding was significant with a p-value of 0.0090. However, when comparing the effect of angiotensin IV (3-8) on spike-ACE2 binding and spike-NRP1 binding, the difference is not significant with a p-value of 0.7296. These results illustrate how angiotensin IV (3-8) has an enhancing effect on spike protein binding to the AXL, ACE2, and NRP1 receptors when compared to the control group. However, the enhancement of spike-AXL binding by angiotensin IV (3-8) is significantly greater than angiotensin IV (3-8)’s enhancement of spike-ACE2 binding and spike-NRP1 binding (Fig. 6D).

### The effects of angiotensin (2-7) and angiotensin (5-7) on spike-AXL binding

To explore the importance of Asp1, Arg2, Val3, and Tyr4 when Phe8 is removed, 40 μM of angiotensin (2-7) and angiotensin (5-7) were applied to the spike-AXL binding assay kits. The addition of angiotensin II (1-8) increased the mean percent absorbance to 130.3%. An unpaired t-test confirmed that adding angiotensin II (1-8) significantly increased spike-AXL binding with a p-value of 0.0380. The addition of angiotensin (2-7) yielded a mean percent absorbance of 195.6%. This angiotensin significantly enhanced spike-AXL binding with a p-value of 0.0014. Additionally, when compared to the angiotensin II (1-8) group, angiotensin (2-7) significantly increased the mean percent absorbance with a p-value of 0.0375. These results suggest that the removal of Asp1 and Phe8 actually increase the enhancing effect of angiotensin II (1-8). Similarly, angiotensin (5-7) had an enhancing effect on spike-AXL binding, concluding a mean percent absorbance of 249.5%. The difference between the untreated group and the angiotensin (5-7) group was statistically significant with a p-value of less than 0.0001. Along with that, the difference between the effect of angiotensin II (1-8) on spike-AXL binding and the effect of angiotensin (5-7) on spike-AXL binding was determined to be significant with a p-value of 0.0002, indicating that the removal of Asp1, Arg2, Val3, Tyr4, and Phe8 do not less angiotensin II (1-8)’s enhancing effect. However, the difference between the angiotensin (2-7) and angiotensin (5-7) groups was not significant with a p-value of 0.1528 (Fig. 7).

**Figure 7:**
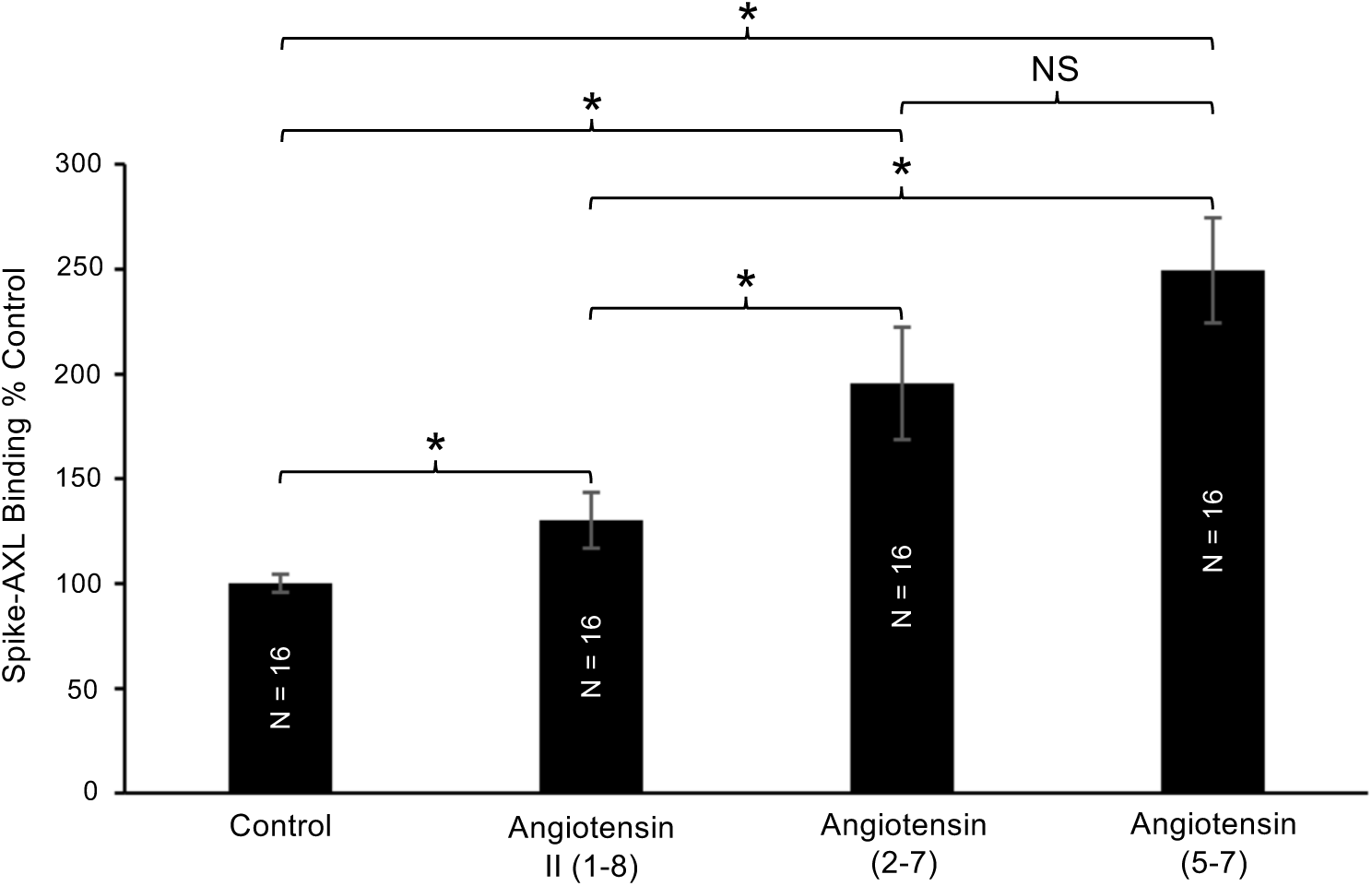
The Effects of Angiotensin II (1-8), Angiotensin (2-7), and Angiotensin (5-7) on Spike-AXL Binding. This figure showcases the effects of angiotensin II (1-8), angiotensin (2-7) and angiotensin (5-7) on spike-AXL binding. Similar to the addition of angiotensin II (1-8) to spike-AXL binding assay kits, the addition of angiotensin (2-7) and angiotensin (5-7) significantly increased spike-AXL binding. Both angiotensin (2-7)’s and angiotensin (5-7)’s enhancing effects on spike-AXL binding were significantly stronger than angiotensin II (1-8)’s enhancing effect on spike-AXL. However, there was not a significant difference between angiotensin (2-7)’s enhancement of spike-AXL binding and angiotensin (5-7)’s enhancement of spike-AXL binding. One asterisk (*) denotes a p-value equal to or less than 0.05 and “NS” represents “not significant”.

### The effects of angiotensin II (1-8) analogs on spike-AXL binding

Wanting to learn more about how essential the amino acids of angiotensin II (1-8) are to the peptide’s overall enhancing effect on spike-AXL binding, angiotensin II (1-8) analogs were applied to the spike-AXL binding assay kits. In this experiment, the addition of angiotensin II (1-8) increased the mean percent absorbance to 130.3%. This difference between the control group’s mean percent absorbance of 100% and the angiotensin II (1-8) group was significant with a p-value of 0.0380 (Fig. 8).

**Figure 8:**
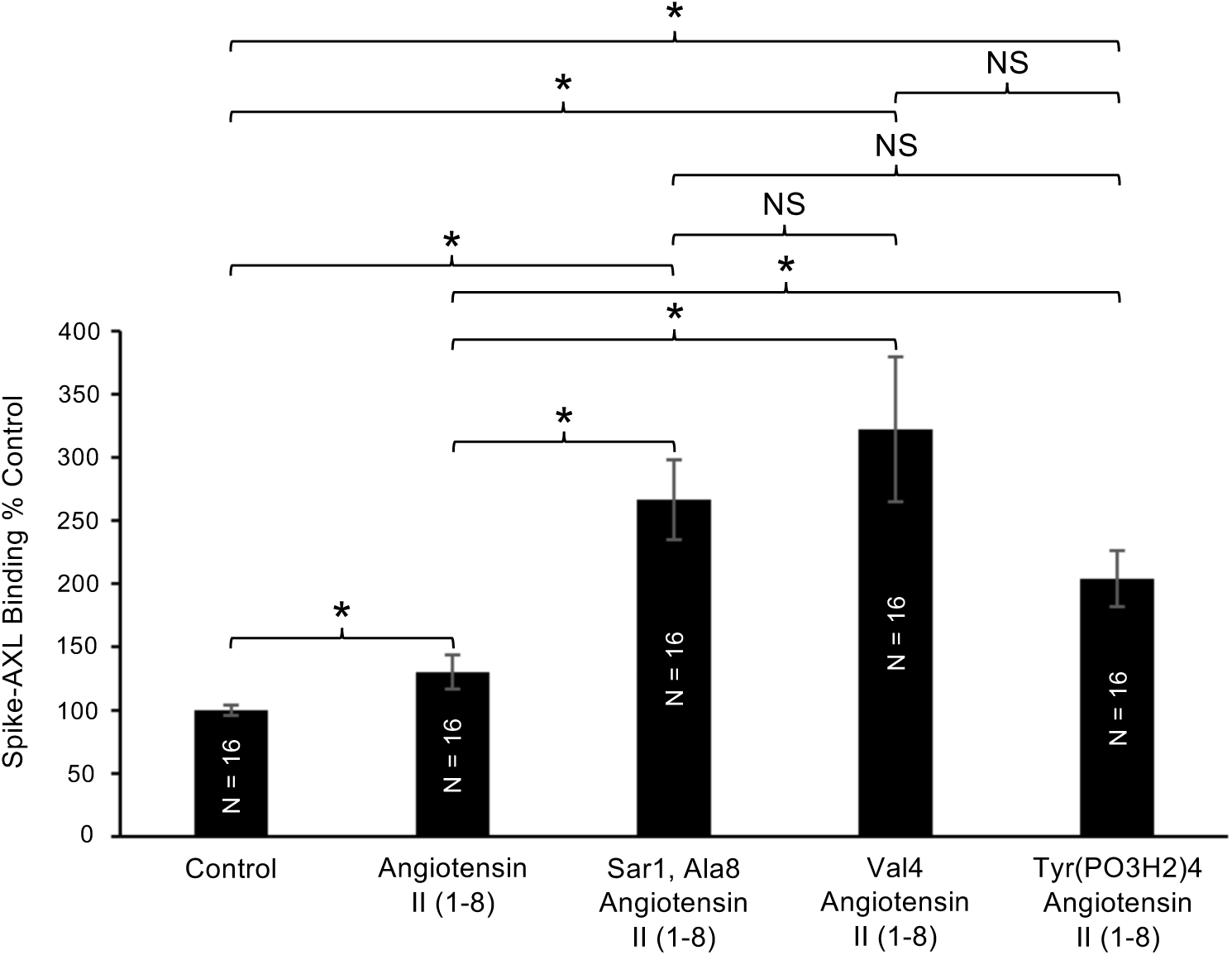
The Effects of Angiotensin II (1-8) and Angiotensin II (1-8) Analogs on Spike-AXL Binding. This figure illustrates the effects of angiotensin II (1-8) and angiotensin II (1-8) analogs on spike-AXL binding. Similar to angiotensin II (1-8), the angiotensin II (1-8) analogs, Sar1, Ala8 angiotensin II (1-8), Val4 angiotensin II (1-8), and Tyr(PO_3_H_2_)4 angiotensin II (1-8), had an enhancing effect on spike-AXL binding. Additionally, the enhancing effects of the analogs on spike-AXL binding were significantly greater than angiotensin II (1-8)’s enhancing effect on spike-AXL binding. However, the differences between the effects of the analogs were not significant. One asterisk (*) denotes a p-value equal to or less than 0.05 and “NS” represents “not significant”.

The Sar1, Ala8 angiotensin II (1-8) analog replaces Asp1 of angiotensin II (1-8) with sarcosine and Phe8 with alanine. The addition of this analog increased the mean percent absorbance to 266.4%. This increased binding interaction was statistically significant with a p-value of less than 0.0001 when compared to the control group’s mean percent absorbance of 100%. Additionally, when compared to the angiotensin II (1-8) group, the Sar1, Ala8 angiotensin II (1-8) analog group’s enhanced spike-AXL binding was significant with a p-value of 0.0004. Despite changing angiotensin II (1-8)’s first and last amino acids, this analog still had an enhancing effect on spike-AXL, suggesting that changing Asp1 and Phe8 will not remove angiotensin II (1-8)’s enhancing effect on spike-AXL binding and, instead, may increase it. This conclusion is confirmed by angiotensin III (2-8) still having an enhancing effect on spike-AXL binding despite lacking Asp1 and angiotensin (1-7) increasing the binding interaction between the spike protein and the AXL receptor even though angiotensin (1-7) is lacking Phe8 (Fig. 8).

The next analog explored was Val4 angiotensin II (1-8), which replaces angiotensin II (1-8)’s Tyr4 with valine. When the Val4 angiotensin II (1-8) analog was applied, the mean percent absorbance was 322.4%. An unpaired t-test found Val4 angiotensin II (1-8)’s enhancing effect to be statistically significant with a p-value of 0.0006 when compared to the control group. In addition, the difference between the effect of angiotensin II (1-8) on spike-AXL binding and the effect of the Val4 angiotensin II (1-8) analog on spike-AXL binding was determined to be significant with a p-value of 0.0028. Although Tyr4 was replaced with valine, this angiotensin II (1-8) analog still increased spike-AXL binding, suggesting that altering Tyr4 does not remove the spike-AXL binding enhancement (Fig. 8).

The last angiotensin II (1-8) analog used was Tyr(PO_3_H_2_)4 angiotensin II (1-8), which has a phosphorylated Tyr4. Similarly, applying this analog yielded a mean percent absorbance of 204.0%. An unpaired t-test confirmed that this analog significantly increased spike-AXL binding with a p-value of less than 0.0001 in comparison to the control group’s mean percent absorbance of 100%. When comparing the angiotensin II (1-8) group and the Tyr(PO_3_H_2_)4 angiotensin II (1-8) analog, the difference was calculated to be significant with a p-value of 0.0080. Despite having a phosphorylated Tyr4, this analog still increased spike-AXL binding, suggesting that this covalent modification on Tyr4 does not change angiotensin II (1-8)’s enhancing effect on spike-AXL binding and, instead, further increases spike-AXL binding. The enhancing effects of the Val4 angiotensin II (1-8) analog and the Tyr(PO_3_H_2_)4 angiotensin II (1-8) analog imply that changing Tyr4 does not influence angiotensin II (1-8)’s enhancing effect on spike-AXL binding. However, when comparing the analogs to one another, their differences in their enhancing effect on spike-AXL binding were not significant (Fig. 8).

## Discussion

SARS-CoV-2, the virus that causes COVID-19 and subsequent PASC, uses its membrane fusion protein, the spike protein, to enter the host cells for infection by binding to receptors [1, 2], such as ACE2 [1, 2, 9, 10], AXL [14], and NRP1 [1]. In addition to the spike protein bound to the viral particles, free circulating spike proteins, in particular, the S1 subunit proteins also bind to these receptors to possibly exert adverse effects that may lead to COVID-19-associated complications. The binding of the spike protein to its major host cell receptor, ACE2, results in the internalization of spike/ACE2 complex, thereby downregulating plasma membrane ACE2 that physiologically functions to degrade angiotensin II (1-8) [21–24]. The reduction of ACE2 levels is expected to increase the levels of angiotensin II (1-8), and indeed plasma levels of angiotensin II (1-8) have been shown to be elevated in COVID-19 patients [25–27]. Increased angiotensin II (1-8) levels would promote AT_1_R-dependent biological actions that may lead to the cardiovascular and neurological complications seen in COVID-19 patients. The major finding of the present study is that increased angiotensin II (1-8) levels would additionally enhance the spike-AXL host cell receptor binding, amplifying the spike protein actions. Another major finding that may even be more significant is that angiotensin IV (3-8), which can be produced from angiotensin II (1-8), potently enhances the spike protein binding to not only AXL but also ACE2 and NRP1.

Caputo et al. [28] reported that angiotensin II (1-8) enhanced the spike protein-dependent infection of wild-type SARS-CoV-2 as well as pseudotyped vesicular stomatitis virus expressing spike protein in Calu-3 human airway epithelial cells. In this study, authors concluded that the mechanism of this event is through angiotensin II (1-8), via AT_1_R, increasing ACE2 mRNA and protein expression [28]. Our results from the present study opens up the possibility that angiotensin II (1-8) may activate the spike protein-AXL binding that, in turn, increases the infection. Alternatively, in their system [28], angiotensin IV (3-8) was produced from the added angiotensin II (1-8) in the culture medium, which then enhanced the spike protein binding to ACE2.

While the direct actions of angiotensin II (1-8) have been well studied, especially through its receptors [3–7], it is not very well known that angiotensin II (1-8) can be cleaved to become other shorter peptides, which can exert other biologic events as consequences of the RAS activation. As shown in Figure 1, these shorter peptides, which include angiotensin III (2-8) and angiotensin IV (3-8), are generated through the actions of aminopeptidases [3–5, 7]. Recently, because of COVID-19, it has become more known that the major spike protein target, ACE2, catalyzes the cleavage of angiotensin II (1-8) to angiotensin (1-7) [3, 4]. While not characterized well, other shorter peptides such as angiotensin (1-6), angiotensin (2-7), and angiotensin (5-7), may also be generated in our body and exert important pathophysiological actions.

Our structure-function study using angiotensin peptides of various lengths as well as analogs indicate that His9 and Leu10 may interfere with the angiotensin peptide and prevent the enhancement of spike-AXL binding as observed in our experiments using angiotensin I (1-10) and angiotensin (1-9). The C-terminal amino acid deletion from angiotensin II (1-8) did not alter the action of this peptide to enhance spike-AXL binding. By contrast, the N-terminal amino acid deletion further enhanced spike-AXL binding, suggesting that the deletion of Asp1, as well as perhaps Arg2, relieved the structural interference that hinders the activation of spike-AXL binding. The results of Sar1, Ala8 angiotensin II (1-8) are consistent with the position that Asp1 interferes with the spike-AXL binding. The results of the experiments involving Val4 angiotensin II (1-8), Tyr(PO_3_H_2_)4 angiotensin II (1-8), and angiotensin (5-7) may also indicate that Tyr4 interferes with the activation of spike-AXL binding, which explains why altering Tyr4 does not remove the peptide’s enhancing effect on spike-AXL binding. Currently, we do not know whether angiotensin peptides interact with the spike protein or the host cell receptors to cause the binding enhancement. However, our results showing that angiotensin IV (3-8) enhances the spike protein binding to all three receptors tested in this study may suggest that the interacting target of angiotensin peptides may be the spike protein.

Although we argued that the spike protein binding to ACE2 is the major factor that increases angiotensin II (1-8) levels, the enhancement of spike protein binding to host cell receptors by angiotensin II (1-8) and shorter angiotensin peptides may also point to an important concept that suggests that hypertensive individuals, who already have higher angiotensin II (1-8) levels, may be more susceptible to SARS-CoV-2 infection, present with more severe symptoms, and be at risk of experiencing further complications.

In summary, it was determined that angiotensin II (1-8) had an enhancing effect on spike-AXL binding, but angiotensin II (1-8) did not significantly modify spike-ACE2 binding or spike-NRP1 binding. The longest length of the angiotensin peptides tested, angiotensin I (1-10), did not have a significant effect on spike-AXL binding. In contrast, shorter lengths of angiotensin peptides, such as angiotensin IV (3-8), notably increased spike-AXL binding. Angiotensin IV (3-8) enhanced the binding of the spike protein not only to AXL, but also to ACE2 and NRP1. Further investigations are needed to determine the roles of angiotensin peptides in enhancing the SARS-CoV-2 infection as well as the actions of freely circulating S1 spike protein, which perhaps promote COVID-19 complications and the symptoms of which patients with long-COVID are currently suffering.

## Acknowledgements

This research was funded by the National Institutes of Health (NIH), grant numbers R21AG073919 and R03AG071596 (to Y.J.S.). The content is solely the responsibility of the authors and does not necessarily represent the official views of the NIH.

